# A computational framework to model cartilage degeneration induced by mechanoinflammation and cytokine-driven inflammation in post-traumatic osteoarthritis

**DOI:** 10.64898/2026.05.29.728618

**Authors:** Moustafa Hamada, Atte S.A. Eskelinen, Joonas P. Kosonen, Sanna Hakonen, Cristina Florea, Alan J. Grodzinsky, Rami K. Korhonen, Petri Tanska

**Author notes:** **Corresponding author at:** Department of Technical Physics, University of Eastern Finland, Yliopistonranta 8, POB 1627, 70211 Kuopio, Finland. Email address, Telephone number: **, **.

## Abstract

Knee joint injury is a major risk factor for post-traumatic osteoarthritis (PTOA), often associated with early cartilage degeneration. Mechanical overloading and cytokine-driven inflammation are key drivers of this process, yet the underlying mechanisms and their distinct temporal and spatial contributions to cartilage degradation remain unclear. Here, we present a mechanobiological finite element framework that simulates cartilage degradation through cell-mediated proteolytic activity triggered by mechanoinflammation and cytokine-driven inflammation. The model reproduces experimentally observed depth-dependent loss of collagen and aggrecan, with mechanoinflammation inducing a transient response and cytokine-driven inflammation sustaining prolonged matrix degradation. Sensitivity analysis further shows that mechanoinflammation-driven degradation is governed mainly by protease production per cell, whereas cytokine-driven degradation is more sensitive to the rate of cellular stimulation. Together, this framework provides a mechanistic basis to study proteolytic cartilage degeneration and supports future *in silico* evaluation of therapeutic strategies aimed at mitigating cartilage degradation in PTOA.

## 1. Introduction

Knee joints support loads several times the body weight while maintaining nearly frictionless motion between articulating surfaces [1]. This performance arises from the unique composition and hierarchical structure of articular cartilage [2,3]. Cartilage is a biphasic tissue consisting of a solid extracellular matrix and an interstitial fluid phase rich in water [2–5]. Its extracellular matrix is formed primarily by a collagen fibril network and highly charged aggrecan proteoglycans, which together enable load support and fluid pressurization [2–5]. Joint injury and the associated upregulation of pro-inflammatory cytokines can disrupt this structural balance [5–7], rendering cartilage vulnerable to loss of tissue homeostasis, in which catabolic activity outpaces anabolic repair[8–10]. When sustained, this imbalance drives irreversible cartilage matrix degeneration and leads to joint dysfunction, impaired mobility, and chronic pain, all of which are key features of post-traumatic osteoarthritis (PTOA) [9,11]. Because PTOA remains a highly prevalent disease without a disease-modifying cure, understanding the cell-mediated mechanisms governing cartilage matrix degeneration is essential for predicting its progression and developing strategies to preserve tissue health.

Decades of experimental work have defined key features of cartilage degeneration across *ex vivo* and *in vivo* models at both tissue and cellular scales [12–16]. Following joint injury, chondrocyte death increases rapidly under elevated tissue strain and stress [17–19]. At the same time, this mechanical overloading drive surviving chondrocytes toward a catabolic state characterized by mitochondrial dysfunction, oxidative stress, and increased production of matrix-degrading proteases, including matrix metalloproteinases (MMPs) such as collagenases MMP-1 and MMP-13), and aggrecanases of the ADAMTS family (a disintegrin and metalloproteinase with thrombospondin motifs), such as ADAMTS-4, and ADAMTS-5 [20,21]. This transmission of mechanical stress and strain to cells through mechanosensitive receptors, followed by activation of catabolic signaling pathways and proteases production is referred to as mechanoinflammation [22]. This phenomenon is associated with cartilage extracellular matrix breakdown, and loss of aggrecan and collagen can begin within days after injury [6,23].

In parallel, cytokine-driven inflammation present after joint injury further amplifies cartilage damage. Pro-inflammatory cytokines, such as interleukin-1 (IL-1) and IL-6, released from the synovium, immune cells, and cartilage itself, diffuse into the tissue and bind to their respective cell surface receptors, activating intracellular signaling pathways that upregulate protease production [24–26]. This signaling can suppress matrix synthesis, increase catabolic protease gene and protein expression, and exacerbate mitochondrial stress and reactive oxygen species production, thereby sustaining the catabolic state and accelerating matrix loss [24–26].

Despite extensive experimental evidence linking both mechanical injury and inflammation to PTOA progression, the cell-mediated biological mechanisms that underline their distinct temporal and spatial contributions to cartilage matrix degradation remain poorly understood. In particular, it remains unclear how proteolytic activity induced by mechanoinflammation and cytokine-driven inflammation independently and jointly contribute to the progression of collagen and aggrecan loss.

Computational modeling provides a powerful tool for investigating the mechanisms linking mechanical overloading and inflammation to cartilage damage under physiologically relevant PTOA conditions [27–31]. Such models can integrate tissue mechanics, cellular responses, and proteolytic processes to provide mechanistic insight into the pathways governing cartilage degeneration [28,31–33]. In turn, they can guide the experimental design, generate new hypotheses, and enable *in silico* testing of therapeutic interventions. However, most existing modeling frameworks focus on either mechanically-driven or inflammation-driven pathways in isolation and often treat the main extracellular matrix components, aggrecan and collagen, separately [34–37]. As a result, there are currently no modelling frameworks that simultaneously resolve the spatial and temporal contributions of mechanoinflammation and cytokine-driven inflammation to proteolytic cartilage degradation.

In this work, we establish a computational framework for simulating temporal and spatial dependent proteolytic cartilage degradation after joint injury and the onset of PTOA due to mechanoinflammation and cytokine-driven inflammation (Figure 1). We hypothesize that proteolytic activity by each of these inflammatory pathways produces distinct temporal and spatial patterns. To test this, the framework couples (1) a biomechanical model that simulates injurious loading and the immediate mechanical response of the tissue, and (2) a biochemical model describing the spatiotemporal degradation of cartilage composition driven by cellular response and proteolytic activity induced by mechanical injurious loading or IL-1 cytokine diffusion. The biochemical processes are described using diffusion–reaction partial differential equations. We evaluated model outcomes against previously published depth-dependent experimental measurements of collagen and aggrecan content at days 0, 3 and 12 post-injury, obtained using Fourier-transform infrared microspectroscopy (FTIR) and digital densitometry (DD), respectively. Because the framework represents detailed cellular responses, it may support future *in silico* evaluation of therapeutic interventions and drug effects.

**Figure 1.**
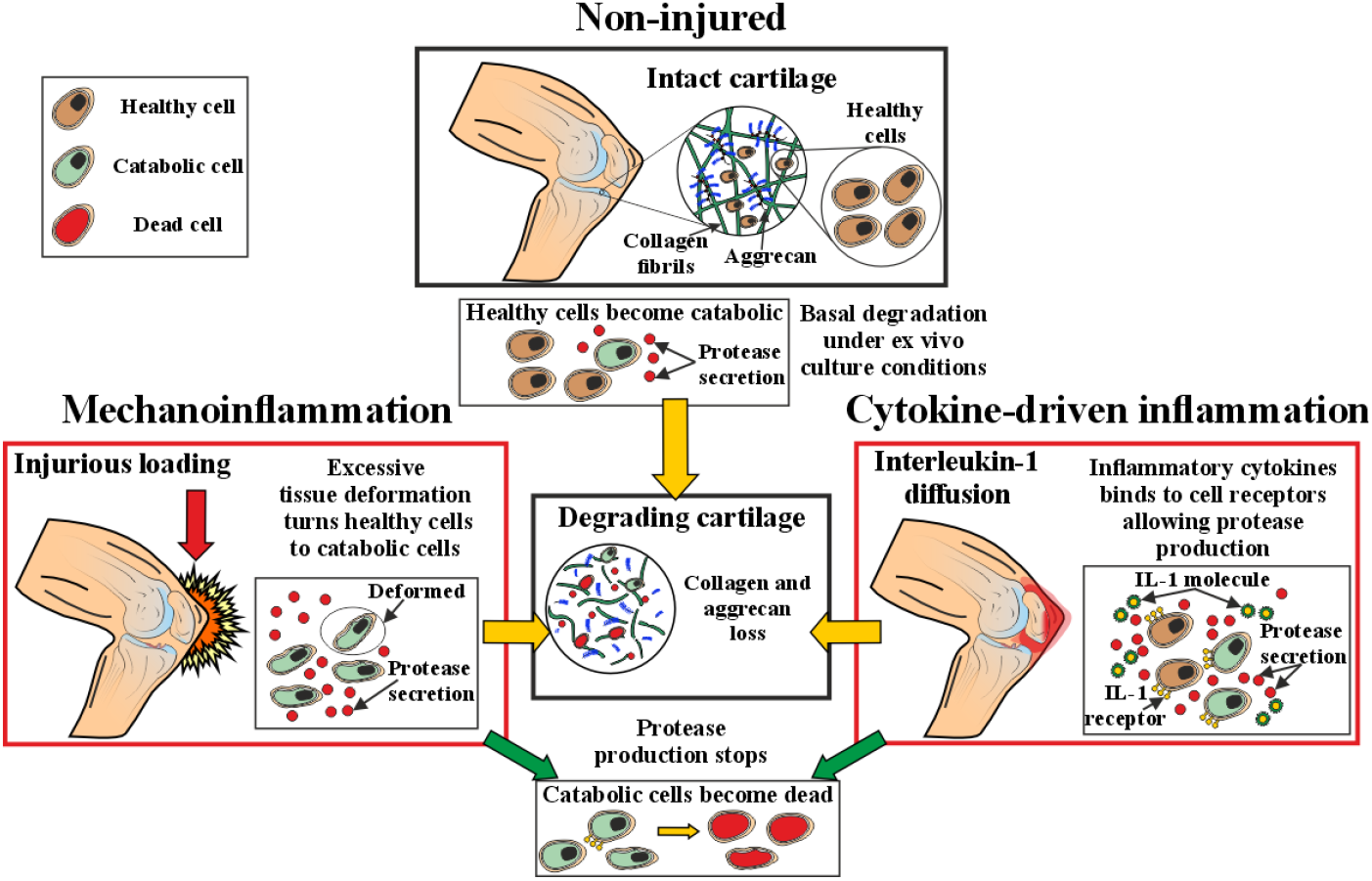
Overview of the cell-mediated mechanisms for collagen and aggrecan loss in articular cartilage following traumatic injury. The model incorporates three cell phenotypes: healthy, catabolic and dead. Matrix-degrading proteases, including MMP and aggrecanase, are produced via two main pathways: (i) mechanoinflammation, which induces the transition of healthy cells into a catabolic state that stimulates proteases production, and (ii) interleukin-1 (IL-1)–driven stimulation of both healthy and catabolic cells via receptor binding. These proteases subsequently degrade collagen fibrils and aggrecan. As catabolic cells transition to a dead state, their protease production ceases. In addition, basal degradation is included to account for low-level proteolytic degradation in the absence of external stimuli reflecting intrinsic limitation of ex vivo culture conditions.

## 2. Methods

### 2.1 Computational modeling framework

In the present work, we aim for the first time to establish computational framework to simulate mechanoinflammation-driven and IL-1-driven inflammatory cartilage degeneration through simultaneous simulation of collagen and aggrecan degeneration (Figure 2A). Model outputs are compared with corresponding *ex vivo* experimental data (Figure 2B) using depth-dependent collagen and aggrecan content measured at days 0, 3 and 12.

**Figure 2.**
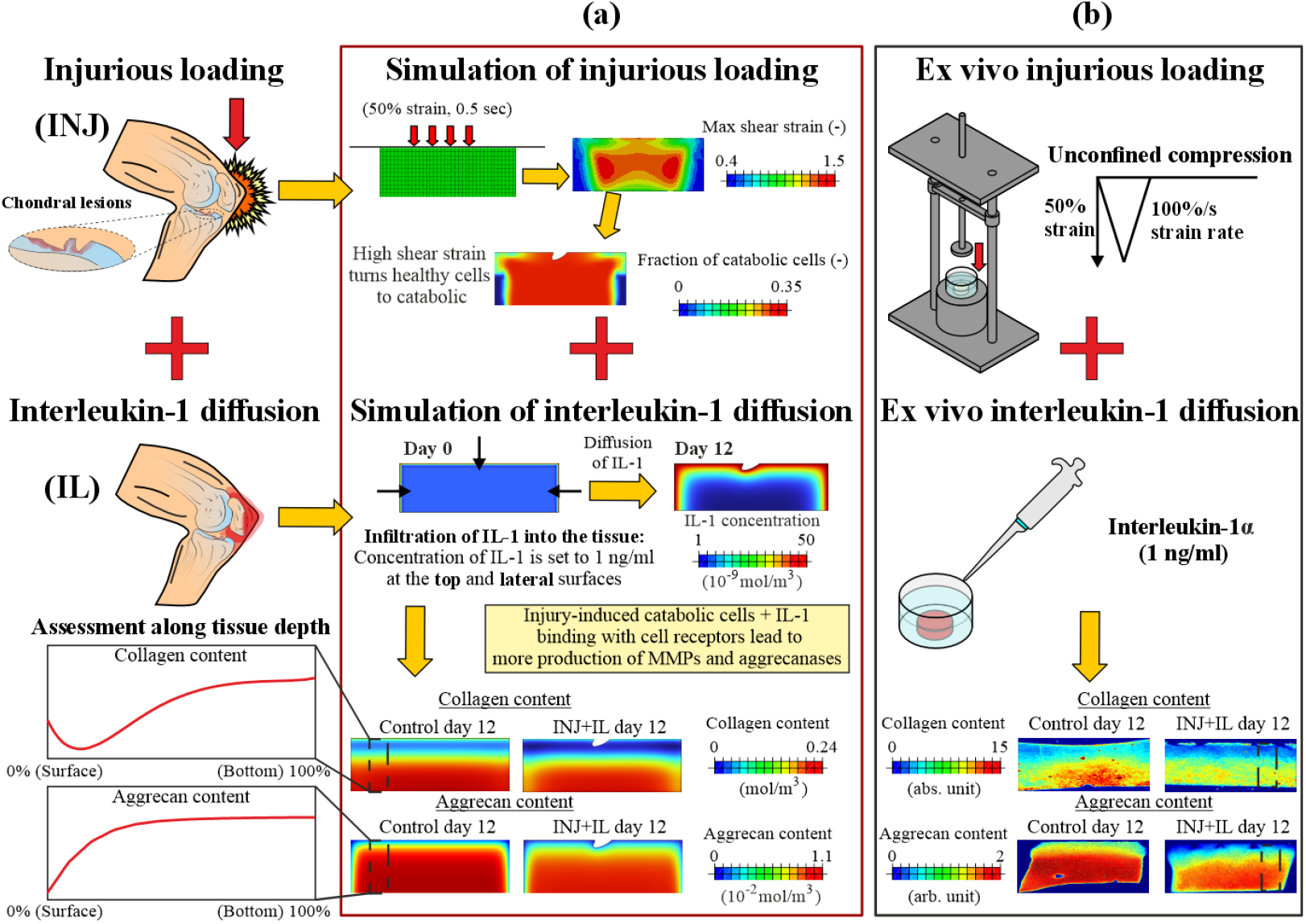
Study workflow for simulating collagen and aggrecan loss in cartilage following traumatic knee injury, integrating injurious loading and interleukin-1 diffusion. A) injurious loading was simulated by applying 50% strain to a 2D cartilage plug (1 mm thickness) at a strain rate of 100%/s. Maximum shear strain was used to estimate the spatial distribution and amount of cell damage, leading to the formation of catabolic cells. In parallel, interleukin-1 (IL-1) infiltration into the tissue was modeled via diffusion over 12 days, stimulating protease production. B) Injurious loading and interleukin-1 diffusion were replicated ex vivo. Cartilage sections were analyzed for collagen content using Fourier transform infrared microspectroscopy and for aggrecan content using digital densitometry. Measurements were obtained at days 0, 3 and 12 post-injury. Model results were compared with experimental findings across tissue depth at the corresponding time points. INJ = Injurious loading, IL = Interleukin-1.

#### 2.1.1 Biomechanical model formulation

To reproduce the injury-driven cartilage degeneration, we first constructed a finite element (FE) model of the *ex vivo* cartilage explant in Abaqus (v. 2023, Dassault Systèmes, Providence, RI, USA), similarly as done previously [38,39]. A single unconfined compression load–unload cycle with 50% compressive strain was applied to an intact 2D cylindrical cartilage geometry (diameter = 3 mm, height = 1 mm) to simulate *ex vivo* tissue injury from the experiments [38,39].

Cartilage was modeled using a user-defined material (UMAT) subroutine that implements a fibril-reinforced porohyperelastic swelling constitutive formulation and captures the depth-dependent composition and complex mechanical behavior of cartilage [29]. Material properties were obtained from experimental data from bovine cartilage explants (see Table S1 in Supplementary Material) [29,40].

#### 2.1.2 Boundary conditions for biomechanical model

Axial displacement of the explant was constrained at the bottom surface of the geometry, while lateral displacement was restricted only at a single node located at the bottom of the symmetry axis, thus allowing lateral expansion of the tissue. Contact between the top surface of the geometry and the rigid compression platen was defined as frictionless surface-to-surface interaction. Fluid flow was permitted through the lateral surfaces by prescribing zero pore pressure.

#### 2.1.3 Biochemical model formulation

A 2D geometry of the cartilage explant was created in COMSOL Multiphysics (version 6.1, MA, USA) using the same dimensions as in the biomechanical model. This biochemical model was designed to simulate cartilage proteolytic responses driven by injurious loading (mechanoinflammation) and IL-1 diffusion (cytokine-driven inflammation) by describing cell catabolism, protease production (MMPs and aggrecanases), and the resulting loss of collagen and aggrecan contents over 12 days. Four model steps were implemented: 1) intact control (CTRL), 2) injury-only (INJ), 3) IL-1 diffusion-only (IL), and 4) combined injury and IL-1 diffusion (INJ+IL).

Time-dependent changes in tissue constituents and cellular population concentrations were described using a system of reaction–diffusion equations of the general form:

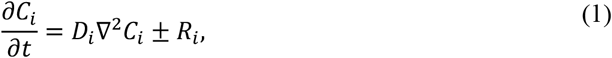

where *C*_*i*_ denotes the concentration of tissue constituent *i, D*_*i*_ is the corresponding effective diffusivity within the tissue, and *R*_*i*_ represents a source or sink term governing production or degradation of constituent *i*.

Cartilage tissue was assumed to comprise three distinct chondrocyte phenotypes representing healthy, catabolic, and dead cellular states. All cell populations were assigned zero diffusivity, based on the assumption that chondrocyte mobility within the cartilage matrix is minimal over 12 days[41]. The temporal evolution of healthy (C_cell,h_), catabolic (*C*_cell,c_), and dead (*C*_cell,d_) cells was governed by:

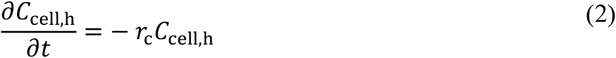

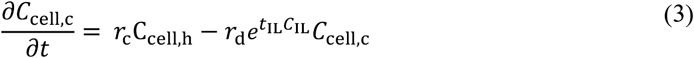

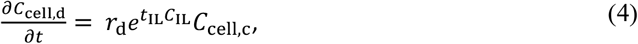

where *r*_c_ and *r*_d_ represent basal transition rates from healthy to catabolic cells and from catabolic to dead cells, respectively [28]. These rates describe transitions between cell populations under *ex vivo* culture conditions in the absence of external stimuli. Their values were calibrated using the CTRL experimental data of depth-dependent collagen and aggrecan content at days 3 and 12, while simultaneously constraining the basal cell death level to remain consistent with previously reported measurements in non-injured cartilage maintained under similar culture conditions (10% of cell death on day 4 of culture) [13,42,43]. These basal cell transitions were the drivers of collagen and aggrecan loss in the CTRL model. The coefficient *t*_IL_ modulates the effect of IL-1 (*C*_IL_) on accelerating the transition of catabolic cells to dead cells in IL-containing models (IL, and INJ+IL) [28]. Healthy and catabolic cells were assumed to be the primary source of protease production in the model; once catabolic cells transitioned to a dead state, protease production was halted.

##### Injury-driven stimulus

In injury-driven models (INJ and INJ+IL), mechanoinflammation was initiated by a fraction of healthy cells transitioning instantaneously into catabolic cells following day-0 injurious loading. Because excessive shear strain has been linked to cell death and catabolic chondrocyte responses [20,44], the spatial distribution of maximum shear strain at 50% compressive strain was obtained from the biomechanical model to define the initial amount and distribution of catabolic cells at day 0. The maximum shear strain (*ε*) was calculated as the maximum difference between the principal components (*ε*_*p*,1_, *ε*_*p*,2_, *ε*_*p*,3_) of the Green–Lagrangian strain tensor [27]:

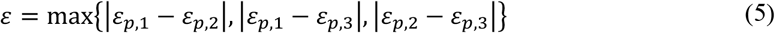

To relate shear strain to injury-driven cell catabolism, a stepwise non-linear function *f*_dmg_(*ε*) was employed [45]:

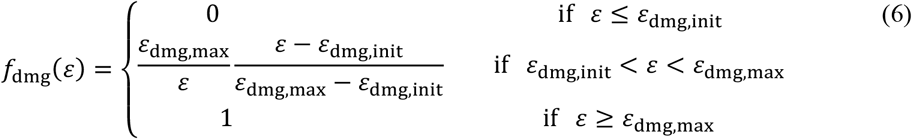

In the regions where the maximum shear strain exceeded 40% (ε_dmg,init_ = 0.4), healthy cells were assumed to transition to the catabolic phenotype at day 0. Above this threshold, the fraction of catabolic cells increases non-linearly with strain, reaching the greatest value at the maximum shear strain of 150% (ε_dmg,max_ = 1.5)[38]. These threshold values are adopted from prior computational studies using fibril-reinforced poroelastic material models and experimental observations of cell apoptosis under excessive strain [38,40,46].

Thereafter, the initial concentration of injury-induced catabolic cells at the start of the biochemical simulation was defined as a fraction of the initial healthy cell concentration:

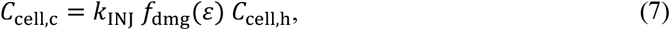

where *k*_INJ_ = 0.3 limits the maximum fraction of injury-induced cell catabolism on day 0 to 30% of healthy cells. This value was based on previously reported measurements of acute cell apoptosis 1 day after injurious loading in bovine cartilage explants [13].

Cells stimulated by injurious loading alter their secretion profiles of MMPs and aggrecanases [47]. To account for delays associated with transcription, translation, and protease secretion, time-dependent stimulus variables *S*_mmp,INJ_ and *S*_aga,INJ_ were introduced [35]. For both the control and injury-driven conditions, the stimulus depended directly on the local concentration of catabolic cells:

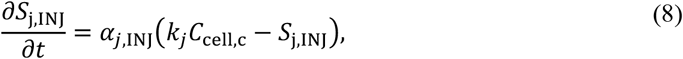

where *j* denotes MMP or aggrecanase, *α*_*j*,INJ_ is stimulus rate constant for *j*, and *k*_*j*_ represents protease production per catabolic cell.

##### Interleukin-1-driven stimulus

In IL simulations (IL and INJ+IL), proteolytic activity was additionally driven by IL-1 diffusion (Figure 2A). Similar to the INJ-driven pathway, MMP and aggrecanase secretions were mediated through time-dependent stimulus variables *S*_mmp,IL_ and *S*_aga,IL_. IL-driven stimulation was governed by IL-1 binding to IL-1 receptors on healthy and catabolic cells:

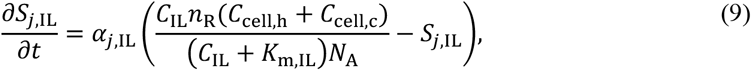

where *n*_R_ is the number of IL-1 receptors per cell, *K*_m,IL_ is a dissociation constant for IL-1–IL-1-receptor binding and *N*_A_ is the Avogadro’s number.

In the combined INJ+IL model, injury- and IL-driven stimuli were both incorporated and treated as independent processes (*S*_*j*,total_ = *S*_*j*,INJ_ + *S*_*j*,IL_).

##### Production of active MMP and aggrecanase

In all models, following cellular stimulations of MMP and aggrecanase secretion, the evolution of active MMP and aggrecanase concentrations, which drive the degradation of intact collagen and aggrecan, respectively, was governed by:

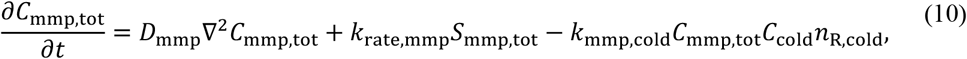

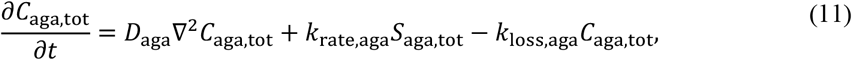

where *k*_rate,mmp_ and *k*_rate,aga_ are rate constants for protease generation, *k*_mmp,cold_ accounts for reduction in the concentration of active MMPs due to binding to degraded collagen (*C*_cold_, Supplementary Material) [35,37], *n*_R,cold_ represents the number of MMP binding sites on degraded collagen, and *k*_loss,aga_ represents rate of aggrecanase degradation [37]. *S*_mmp,tot_ and *S*_aga,tot_ are the total stimulus variables. In the INJ-only and IL-only models, the total stimulus is *S*_*j*,total_ = *S*_*j*,INJ_, and *S*_*j*,total_ = *S*_*j*,IL_, respectively.

Degraded collagen (*C*_cold_) and aggrecan (*C*_aggd_) were defined as separate species as their accumulation modulate protease binding kinetics and thereby indirectly influence matrix net proteolytic degradation [35]. The governing equations for degraded collagen and aggrecan, along with all parameter values used in the biochemical model, are provided in the Supplementary Material. For simplicity, MMPs and aggrecanases were treated as aggregate species representing multiple degrading proteases (e.g., MMP-1 and MMP-13 for MMPs; ADAMTS-4 and ADAMTS-5 for aggrecanases).

##### Collagen and aggrecan loss

The change in collagen (*C*_col_) and aggrecan (*C*_agg_) contents over time across all modelling scenarios were described as:

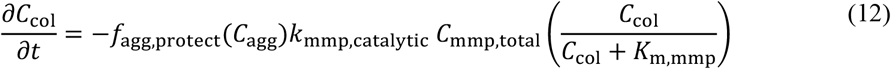

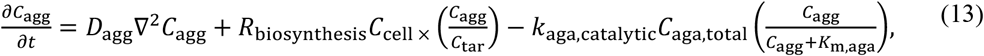

Collagen loss was driven by proteolytic degradation of MMPs while modulated by the local aggrecan content, described with *f*_agg,protect_(*C*_agg_). This represents hindered access of MMPs to intact collagen fibrils due to presence of aggrecan (see Supplementary Material)[35,48]. The parameter *k*_mmp,catalytic_ defines the catalytic activity rate of MMPs, while *K*_m,mmp_ is the Michaelis constant describing the MMP concentration at which collagen cleavage proceeds at half-maximum rate [35]. We did not consider collagen biosynthesis in the model, as the turnover of collagen in cartilage is significantly low compared to aggrecan and for this early stage of injury collagen biosynthesis might be negligible [16,49,50].

Aggrecan loss was governed by diffusion, biosynthesis, and proteolytic degradation of ADAMTS. The biosynthesis term *R*_biosynthesis_ represents depth-dependent aggrecan production in the tissue (see Supplementary Material) and is regulated toward a target homeostatic concentration *C*_tar_ above which aggrecan synthesis ceases. Aggrecan degradation was mediated by aggrecanases, with *k*_aga,catalytic_ defining the catalytic activity rate and *K*_m,aga_ representing the Michaelis constant for aggrecanases binding to intact aggrecan.

#### 2.1.4 Boundary conditions for the biochemical model

Zero flux boundary conditions were prescribed at the bottom surface of the cartilage explant, reflecting the restriction of mass transport across this surface. At the top and lateral surface, Robin boundary conditions were applied for intact aggrecan, degraded aggrecan, and MMP by defining concentration gradient perpendicular to these tissue surfaces to be a function of the change of concentration across cartilage–culture medium surface [35]. Dirichlet boundary conditions (value = 0) were imposed for degraded collagen and aggrecanases at these same surfaces, representing low amount of these molecular species in the culture medium [35].

The initial healthy cell concentration was assumed to be spatially uniform throughout the tissue, consistent with immature bovine articular cartilage [35]. Initial concentrations of MMPs, aggrecanases, degraded collagen and degraded aggrecan were set to zero.

The initial depth-dependent collagen concentration was defined by scaling the collagen content obtained from the FTIR measurements (*C*_col,FTIR_), expressed in absorption unit to the mean collagen concentration of 0.2 moles/m^3^ as reported by Kar et al[35]:

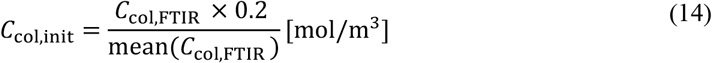

Similarly, the initial aggrecan distribution was determined from the fixed charge density (FCD, *C*_agg,FCD_) expressed in mEq/ml as reported in Orozco et al. [40]. FCD was estimated as two moles of negative charge per mole of chondroitin sulfate disaccharide (CSD), the fundamental repeating unit of chondroitin sulfate and a major component of aggrecan. To convert FCD values to aggrecan concentration (mol/m^3^), the measurements were scaled using the molecular weight of the CSD (502.5 mg/mol) as follows:

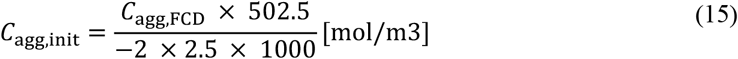

#### 2.1.5 Sensitivity and model results analysis

All model outputs are presented as depth-dependent profiles and compared against experimental findings for collagen and aggrecan content on day 3 and day 12. The model–experiment agreement was assessed based on whether model depth-wise profile outputs fell within the experimental 95% confidence intervals (CIs), reflecting consistency with observed biological and experimental variability.

In addition, to quantify the distinct temporal contributions of each inflammatory pathway to protease activity in INJ+IL model, we estimated the ratios of mechanoinflammation-driven 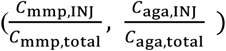 and IL-1-driven 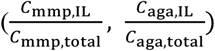 active protease concentrations to their total respective concentrations over time.

A sensitivity analysis was conducted to assess whether mechanoinflammation-driven matrix loss is more strongly governed by the amount of protease produced per catabolic cell (*k*_mmp_, and *k*_aga_) or by the rate of protease stimulus (*α*_mmp,INJ_ and *α*_aga,INJ_). Additionally, we assessed whether the rates of protease stimulus by cytokine-driven inflammation (*α*_mmp,IL_ and *α*_aga,IL_) make a significant contribution to cartilage degradation. All parameters are varied by ±50% from the reference values. For the INJ+IL model, sensitivity analysis was also conducted on same parameters governing the rate of protease stimulus (*α*_mmp,INJ_, *α*_mmp,IL_ *α*_aga,INJ_, and *α*_aga,IL_) to test how they affect the distinct temporal contributions of mechanoinflammation and cytokine-driven inflammation to total active MMP and aggrecanase concentrations.

### 2.2 Experimental work

#### 2.2.1 Sample preparation

All experimental work described here has been performed and published previously and was used here for calibration and comparison with model outputs [18,19]. Cylindrical cartilage explants (*n* = 60; diameter = 3 mm, height = 1 mm) were harvested from four locations within the patellofemoral grooves of knee joints obtained from 1- to 2-week-old bovines (*N* = 8) [18,19]. Prior to the experimental protocols, explants were equilibrated in serum-free culture medium for two days. During the experimental period, the culture medium was changed every two days [19].

#### 2.2.2 Experimental cartilage injury and inflammation protocols

To mimic cartilage degeneration following traumatic injuries under controlled laboratory conditions, an *ex vivo* cartilage model of early-stage PTOA was employed [15,50,51]. Cartilage explants were assigned to one of four experimental groups: (1) injurious loading (INJ), (2) IL-1α-challenge (IL), (3) combined injurious loading and IL-1α-challenge (INJ+IL), or (4) free-swelling, non-injured control (CTRL). For each experimental group, culture was terminated at days 3 or 12 (*n* = 6 plugs per group per time point). Culture was also terminated for the INJ and CTRL groups only at day 0.

The injurious loading protocol was applied by subjecting cartilage explants to a single load-unload cycle of unconfined compression (50% strain amplitude, 100%/s strain rate), using a custom-designed incubator-housed loading device (Figure 2B) [44]. To mimic exogenous inflammation, 1 ng/ml of pro-inflammatory cytokine IL-1α was added to the culture medium, allowing cytokine diffusion into the tissue. The selected concentration was based on prior dose-response tests [19]. Both protocols have previously been shown to cause bulk and localized loss of collagen and aggrecan [18,50].

#### 2.2.3 Collagen and aggrecan content quantification

Collagen content was estimated using Fourier-transform infrared microspectroscopy (FTIR; Agilent Cary 670/620; Agilent Technologies, Santa Clara, CA, USA; pixel size 5.5 µm, spectral resolution 4 cm^-1^; Figure 3A). For each explant, three unstained tissue sections (5-µm-thick) were prepared. The area under the Amide I absorption band (wavenumber range: 1580 cm^-1^ to 1720 cm^-1^) was calculated for each section, and pixel-wise Amide I absorption intensity area was used as an indicator of collagen content [52,53].

**Figure 3.**
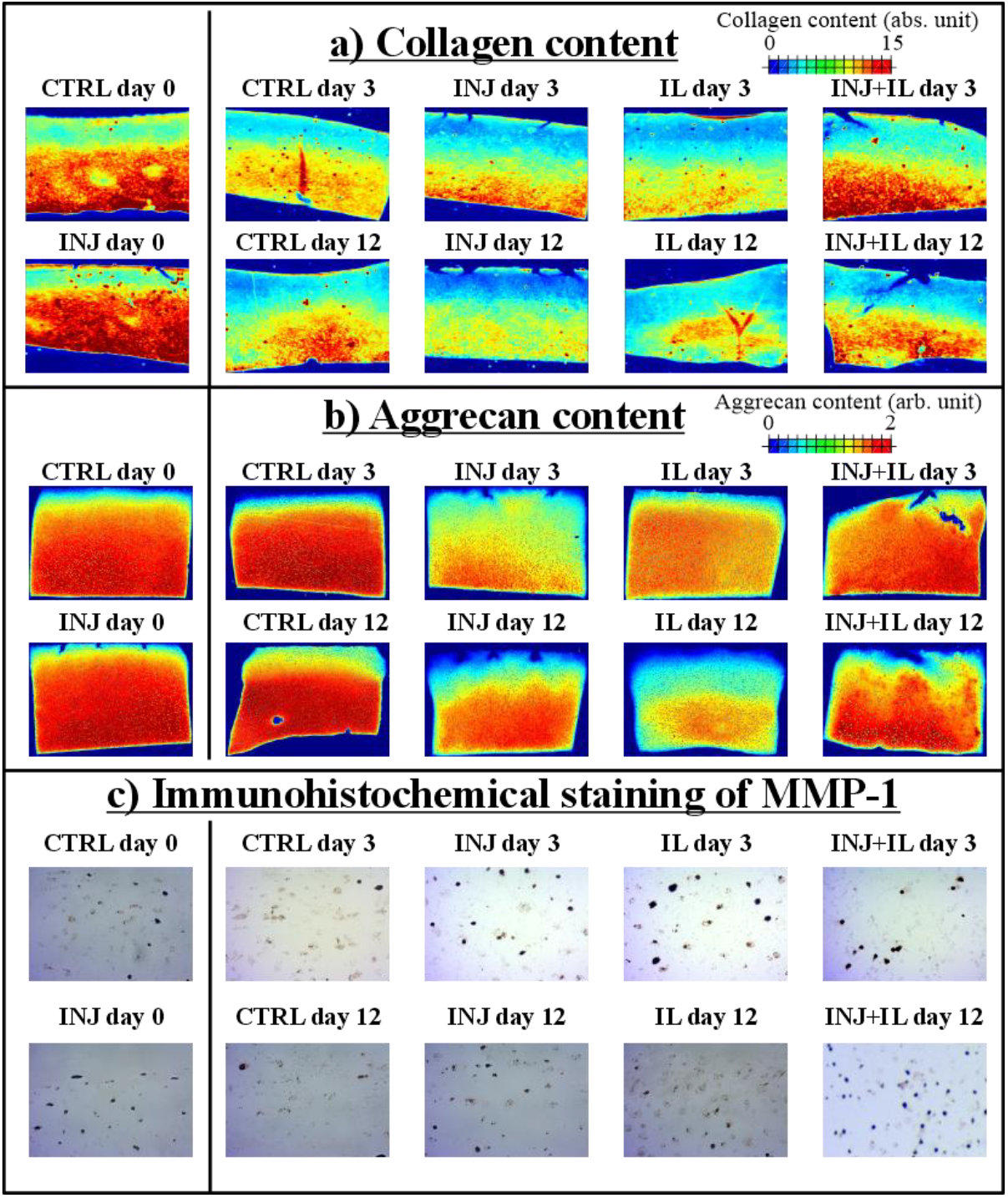
Experimental assessment of cartilage matrix composition and immunohistochemical staining following injurious loading and interleukin-1 diffusion. Histological sections obtained from cartilage explants were used to assess A) collagen content with Fourier-transform infrared microspectroscopy, B) aggrecan content via optical density of Safranin-O-stained sections using digital densitometry and C) immunohistochemical staining of MMP-1, reflecting potential protease activity within the tissue. Chemical maps show lower collagen and aggrecan content in INJ, IL, and INJ+IL groups compared to CTRL on day 12. Immunohistochemical analysis further reveals qualitatively higher staining of MMP-1 in INJ, IL, INJ+IL groups relative to CTRL on days 3 and 12. CTRL = non-injured control, INJ = injurious loading, IL = interleukin-1, MMP = matrix-metalloproteinases.

Aggrecan content was assessed (Figure 3B) using digital densitometry (DD) with a conventional light microscope (Nikon Microphot FXA, Nikon Inc., Tokyo, Japan; 4× magnification, pixel size 1.23 µm). For each explant, three Safranin-O-stained sections (3-µm-thick) were analyzed, and optical density values were used as a measure of aggrecan content [19].

To characterize depth-wise distributions of collagen and aggrecan, two full-thickness regions of interest (ROI; width 200 µm) were defined on the left and right sides of each FTIR and DD chemical maps [18,19]. Values extracted from the ROIs were averaged laterally and normalized by tissue depth (0% = surface, 100% = bottom) to obtain a depth-wise profile of collagen and aggrecan content from each section per explant.

#### 2.2.4 Immunohistochemical staining

MMP-1 IHC staining was performed to qualitatively assess protease upregulation in cartilage at different injury conditions (Figure 3C). For each experimental group, two 4 µm-thick sections were analyzed. An antibody specific for bovine tissues was used (Biomatik, Ontario, Canada). Imaging of the stained sections was performed using a light microscope (Zeiss Primostar 3, Jena, Germany) with 40× magnification. More details on the protocol used for the staining can be found from our previous work [18].

#### 2.2.5 Statistical analysis

All statistical analyses of experimental data were performed using linear mixed-effects (LME) models implemented in IBM SPSS Statistics 29.0 (IBM company, Armonk, NY, USA). LME models were employed to account for the hierarchical structure of the data, as explants were harvested from multiple anatomical locations within the same animal. In these statistical models, animals were identified as the subject with random intercepts to account for inter-animal biological variability. A diagonal covariance was used to accommodate unequal numbers of explants across harvesting locations and experimental groups. Experimental group was included as a fixed factor. Experimental results are presented as LME model estimated mean depth-wise collagen and aggrecan contents with ±95% confidence intervals. When comparing depth-wise profiles between two experimental groups, differences at a given normalized depth were considered statistically significant (*p* < 0.05) if the mean value of one group lay outside the ±95% confidence intervals of the other group.

## 3. Results

### 3.1 Model vs. experimental results: Collagen content

On day 0, no significant differences between the CTRL and INJ groups were observed along normalized tissue depth, neither in the experiment nor in the model outputs (Figure 4A).

**Figure 4.**
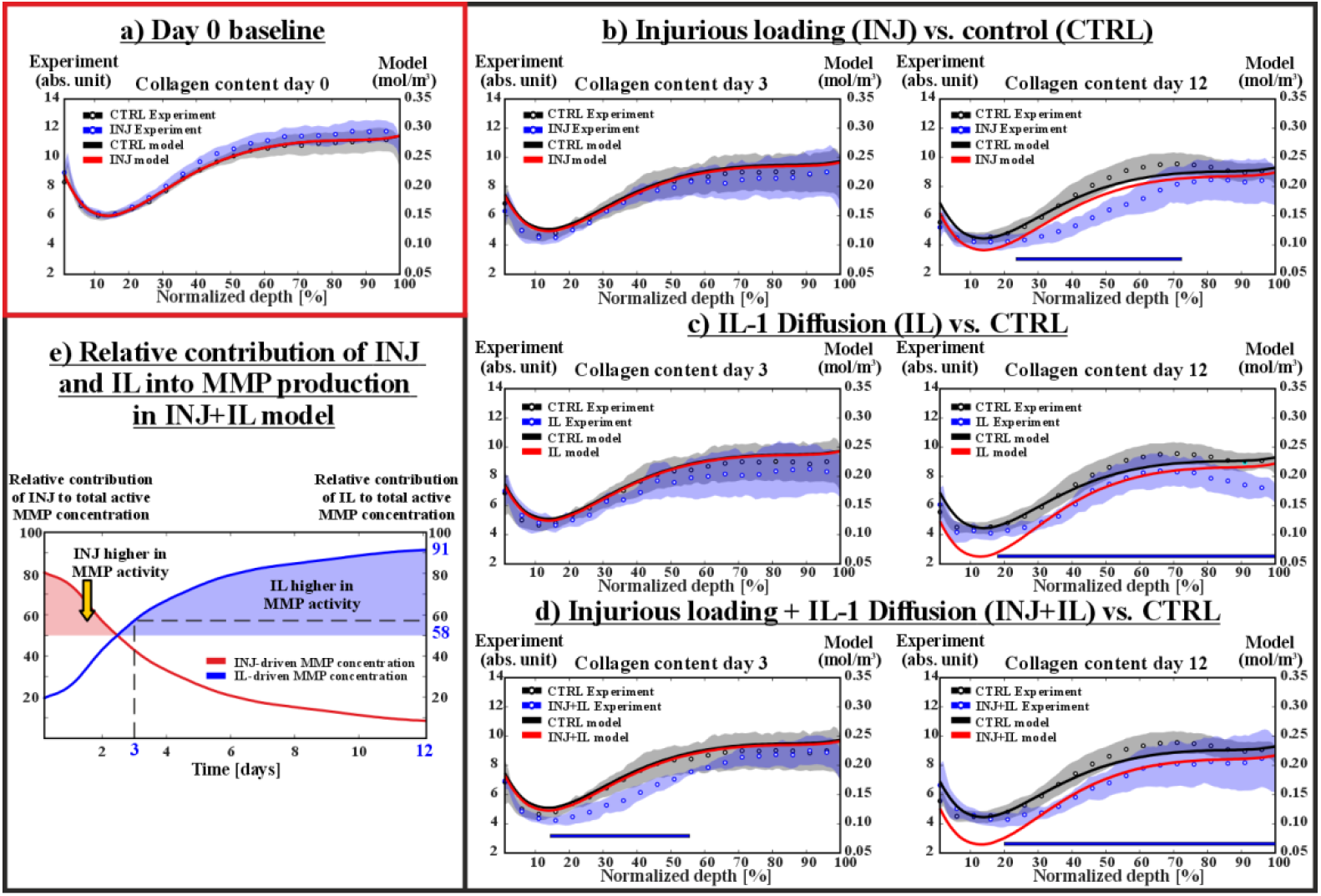
**Comparison between model results and experimental measurements of collagen content** for A) baseline comparison on day 0, between injurious loading group to control. B-D) Comparisons on days 3 and 12 showing collagen content in response to injurious loading, interleukin-1 diffusion, and their combination relative to control. E) Simulated relative contribution of injurious loading and IL-1 diffusion to the total MMP concentration driving collagen degradation over time. Model depth-wise profiles on day 12 fall within the 95% confidence interval of experimental data across most tissue depth for INJ, IL, and INJ+IL when compared to CTRL. IL-1 accounted for approximately 58% and 91% of the total MMP concentration on days 3 and 12, respectively. The blue bar in (B-D) indicate regions across tissue depth with statistically significant differences between experimental groups. CTRL = non-injured control, INJ = injurious loading, IL = interleukin-1, MMP = matrix-metalloproteinases.

On day 3, experimental depth-wise profiles of collagen content showed no significant differences between the INJ or IL groups and CTRL across the full tissue thickness (Figure 4B&C). Consistent with these findings, the simulated INJ profile remained within the experimental 95% CI of the INJ and IL groups across most of the tissue depth. In contrast, the combined INJ+IL experimental group exhibited a significant reduction in collagen content compared to CTRL, with an average decrease of 17% within 15–56% of normalized depth (Figure 4D). The simulated INJ+IL group showed a minor decrease in collagen content of 3% primarily within 0– 20% of the tissue. Agreement between the INJ+IL model and the corresponding experimental group was observed in the deep region (80–100% depth), where the model profile fell within the experimental 95% CI.

On day 12, experimental measurements revealed a pronounced reduction in collagen content across all injured or inflamed groups compared to CTRL. In the INJ group, collagen content was significantly reduced by 24% on average at 23–74% of normalized depth compared to CTRL (Figure 4B). The corresponding model profile fell within the experimental 95% CI in the superficial (0–30%) and deep (66–100%) regions, capturing an average collagen content reduction of 15%.

For the IL group, experiments showed a significant decrease in collagen content of 15% on average across a broad depth range of 18–100% (Figure 4C). The corresponding model profile aligned with the experimental 95% CI within 28–88% of the tissue depth, showing an average collagen reduction of 11%.

The INJ+IL group exhibited a significant reduction in collagen content of 18% on average within 20–100% depth (Figure 4D). The INJ+IL model profile was within the experimental 95% CI over 31–100% depth, capturing an average reduction in collagen content of 15% relative to the CTRL model.

### 3.2 Model vs. experimental results: Aggrecan content

On day 0, no statistically significant differences were found in aggrecan content between the INJ and CTRL groups throughout tissue depth. Similarly, no substantial differences were observed between the simulated INJ and CTRL profiles (Figure 5A).

**Figure 5.**
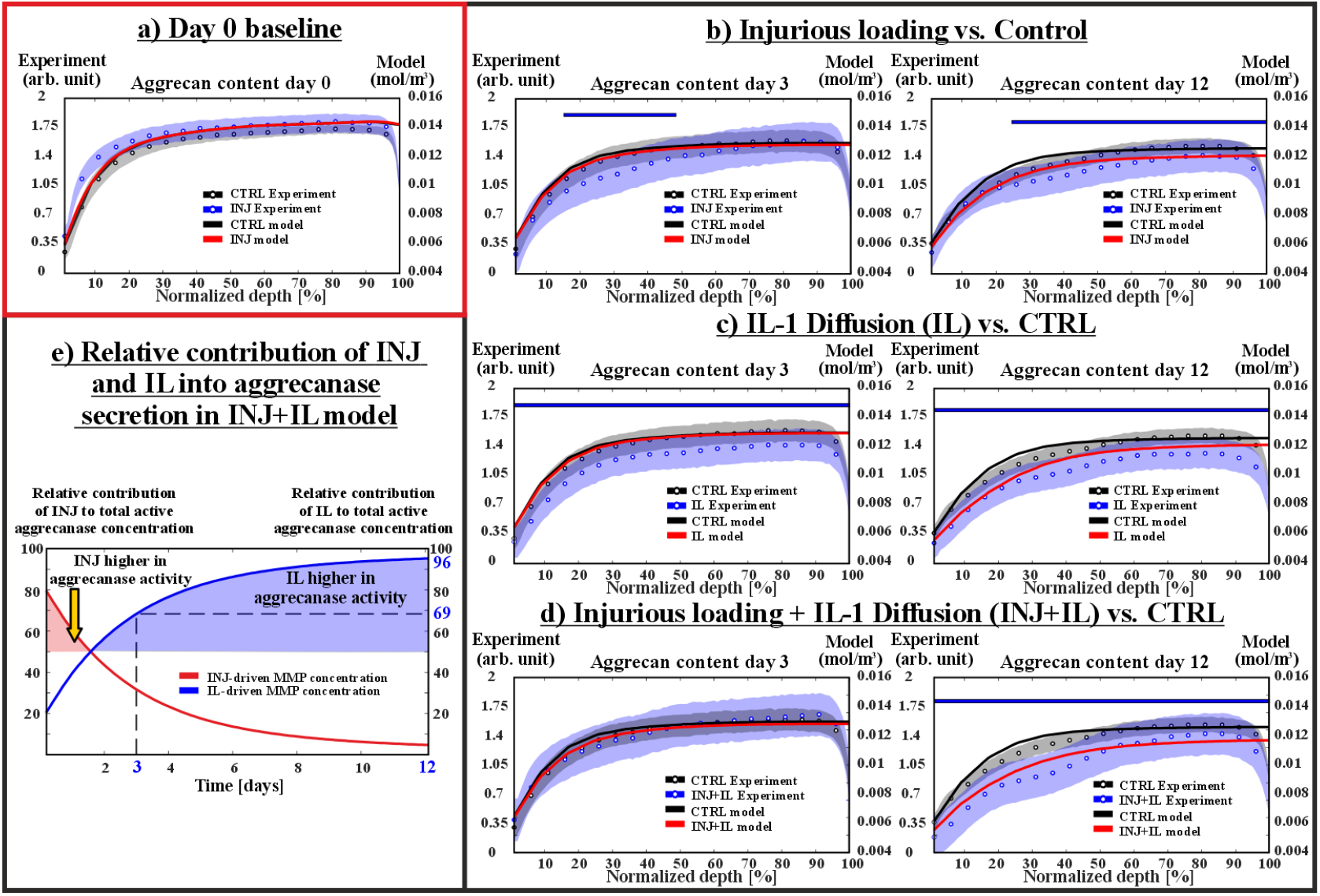
**Comparison between model results and experimental measurements of aggrecan content for** A) baseline comparison on day 0, between injurious loading group to control. B-D) Comparisons on days 3 and 12 showing aggrecan content in response to injurious loading, interleukin-1 (IL-1) diffusion, and their combination relative to control. E) Simulated relative contribution of injurious loading and IL-1 diffusion to the total aggrecanase concentration driving aggrecan loss over time. Model depth-wise profiles on day 12 fall within the 95% confidence interval of experimental data across most of tissue depth for INJ, IL, and INJ+IL when compared to CTRL. IL-1 accounted for approximately 69% and 96% of the total aggrecanase concentration on days 3 and 12, respectively. The blue bar in (B-D) indicate regions across tissue depth with statistically significant differences between experimental groups. CTRL = non-injured control, INJ = injurious loading, IL = interleukin-1, MMP = matrix-metalloproteinases.

On day 3, experimental measurements for the INJ group revealed a significant reduction in aggrecan content relative to CTRL, with an average decrease of 13% within 16–47% depth (Figure 5B). The corresponding INJ model profile showed an average aggrecan decrease of 3% within 10–30% depth. However, the model profile remained within the experimental 95% CI across the full tissue depth. For the IL group, experimental aggrecan content was significantly lower than CTRL throughout the entire tissue depth, with an average reduction of 13% (Figure 5C). The IL model profile showed an average decrease in aggrecan content of 3%, while remaining within the 95% CI across all depths. For the INJ+IL group, no significant differences in aggrecan content were observed experimentally compared to CTRL (Figure 5D). The INJ+IL model profile overlapped with the experimental profiles and lay within the corresponding 95% CI across the full tissue depth.

On day 12, experimental measurements showed a significant reduction in aggrecan content in all injured or inflamed groups compared to CTRL. In the INJ group, aggrecan content was on average 10% lower within 25–100% depth (Figure 5B). The simulated profile captured this trend and remained within the experimental 95% CI across the full tissue depth, with an average aggrecan content reduction of 10%.

For the IL group, experimental measurements showed a pronounced decrease in aggrecan content of 17% across the entire depth (Figure 5C). The corresponding model profile aligned with the experimental profiles within the 95% CI, showing an average decrease in aggrecan content of 12%.

Similarly, the INJ+IL group experimental measurements showed a pronounced decrease in aggrecan content of approximately 20% throughout the tissue depth (Figure 5D). INJ+IL model profile was in agreement with experimental variability, remaining within the 95% CI and showing an average aggrecan content decrease of 25%.

### 3.3 Relative temporal contributions of mechanoinflammation and IL-1-driven pathways to active protease concentrations

In the combined INJ+IL model, the relative contributions of the mechanoinflammation and IL-1-driven pathways to the total matrix-targeting protease concentrations were quantified over time to determine how each pathway contributed to proteolytic activity.

For MMPs, the mechanoinflammatory pathway dominated up to day 2 post-injury, accounting for 60% of the total active MMP concentration at this time point (Figure 4E). This contribution then declined rapidly, and by day 3, IL-1 signaling became the main driver, accounting for 58% of the total active MMP concentration. The IL-1-driven contribution continued to increase thereafter, reaching 91% of the total active MMP concentration by day 12.

For aggrecanase, the mechanoinflammatory pathway was the dominant contributor only during the first day after injury, accounting for 58% of the total active aggrecanase concentration (Figure 5E). By day 3, IL-1 signaling became the primary contributor, accounting for 69% of the total active aggrecanase concentration, and this contribution further increased to 96% by day 12.

### 3.4 Sensitivity analysis

To determine whether mechanoinflammation-driven matrix loss in the INJ model was more strongly governed by the amount of protease produced per catabolic cell or by the rate of catabolic cells protease stimulation, sensitivity analyses were performed on the parameters k_mmp_, k_aga_, α_mmp,INJ_, and α_aga,INJ_ in the INJ model. Increasing the amount of MMP produced per catabolic cell from the reference value (k_mmp_ = 2.5 × 10^−22^ [mol]) to 3.75 × 10^−22^ [mol] resulted in 6% greater collagen loss by day 12 (Figure 6A). Similarly, increasing the aggrecanase production per cell (k_aga_) by the same factor from the reference value resulted in 8% greater aggrecan loss (Figure 6B). In contrast, increasing the protease stimulus rates (α_mmp,INJ_, α_aga,INJ_) from the reference value to 6 × 10^−6^ [s^−1^] resulted in only minor additional loss (~1%) in collagen and aggrecan content (Figure 6C&D).

**Figure 6.**
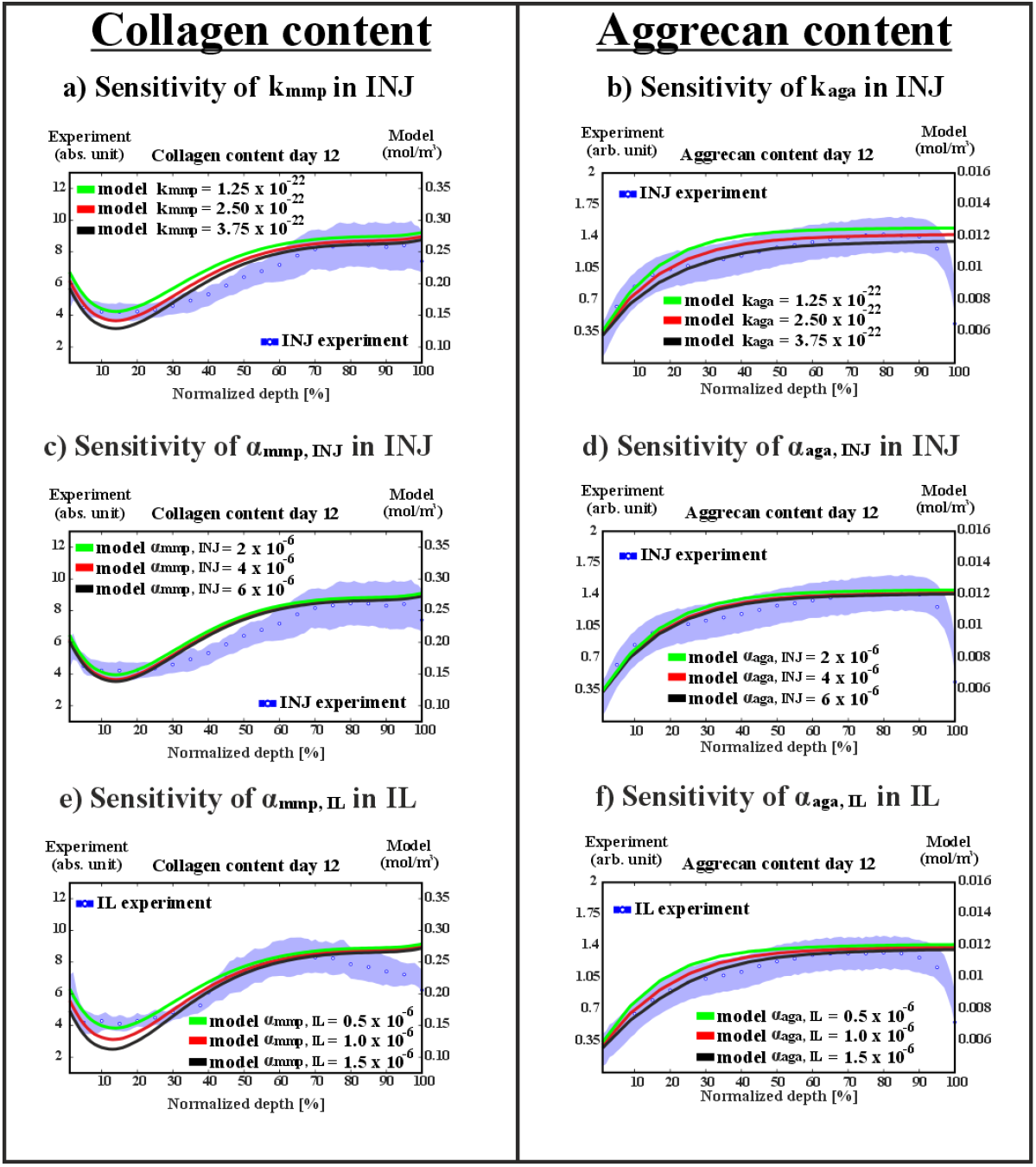
Effect of selected parameters on depth-wise collagen and aggrecan loss. A-B) The amount of MMP and aggrecanase produced per cell, C-D) rates of MMP and aggrecanase stimulus by injurious loading, E-F) rates of MMP and aggrecanase stimulus by interleukin-1 diffusion. INJ = injurious loading, IL = interleukin-1, MMP = matrix-metalloproteinases.

On the other hand, increasing the inflammatory IL-1 stimulus rates (α_mmp,IL_, α_aga,IL_) from the reference value of 1 × 10^−7^ [s^−1^] to 1.5 × 10^−7^ [s^−1^] resulted in 6% greater loss of both collagen and aggrecan contents (Figure 6E&F).

Finally, sensitivity analysis done for the combined INJ+IL model revealed that increasing α_mmp,INJ_ and α_aga,INJ_ to 6 × 10^−6^ [s^−1^] had negligible effects on temporal contributions of the mechanoinflammation and IL-1-driven pathways. Specifically, the mechanoinflammatory contribution increased only from 41% to 42% of total active MMP concentration on day 2 (Figure 7A) and from 31% to 32% of total aggrecanase concentration at the same time point (Figure 7B). When α_mmp,IL_ and α_aga,IL_ increased to 1.5 × 10^−7^ [s^−1^], the INJ contribution to total MMP concentration was reduced from 57% to 45% on day 2, and the contribution to total aggrecanase concentration was reduced from 43% to 34%. IL contribution increased from 58% to 68% of the total active MMP concentration on day 3 (Figure 7C) and from 69% to 77% of the total aggrecanase concentration (Figure 7D). By day 12, IL contributed 94% of total MMP concentration compared to 91% and contributed 97% of total aggrecanase concentration compared to 95% in the reference model.

**Figure 7.**
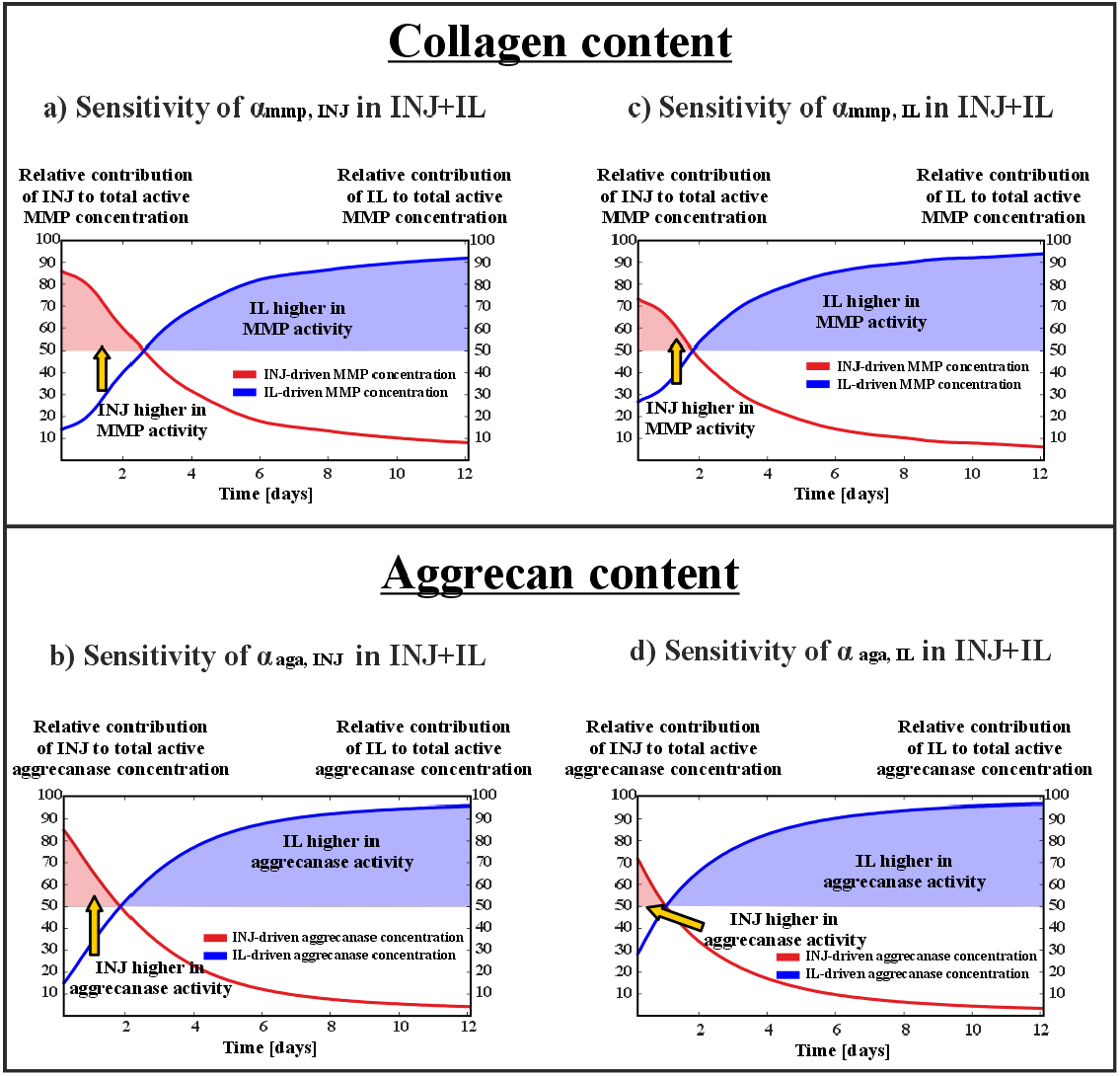
Effect of selected parameters on the relative contribution of injurious loading and interleukin-1 diffusion to MMP and aggrecanase activity. A-B) rates of MMP and aggrecanase stimulus by injurious loading, C-D) rates of MMP and aggrecanase stimulus by interleukin-1 diffusion. INJ = injurious loading, IL = interleukin-1, MMP = matrix-metalloproteinases.

## 4. Discussion

We developed a time-dependent computational framework to simulate how mechanoinflammation (injurious loading) and cytokine-driven inflammation (IL-1 diffusion) differentially drive cell-mediated proteolytic degradation of collagen and aggrecan. In the model, protease activity is promoted by injury-induced catabolic cells and by IL-1 driven stimulation of healthy and catabolic cells. Three main conclusions emerge from our simulations. First, IL-1-driven proteolytic activity represents depth-dependent decrease in collagen and aggrecan content observed experimentally after 12 days, whereas mechanoinflammation-driven proteolytic activity accounts for the depth-dependent aggrecan loss but only partially explains the collagen loss (Figure 4B). Second, when injurious loading and IL-1 diffusion are combined, mechanoinflammation produces an acute short-term proteolytic response, whereas IL-1-driven inflammation sustains longer-term matrix degradation, indicating distinct temporal windows for the two pathways. Third, mechanoinflammation-driven cartilage degradation is influenced more strongly by the amount of protease produced per catabolic cell (parameters k_mmp_, and k_aga_) than by the rate at which cells are stimulated to produce proteases (parameters α_mmp,INJ_, and α_aga,INJ_). In contrast, when proteolytic activity is driven by IL-1-inflammatory signaling, the rate of protease stimulation (parameters α_mmp,IL_, and α_aga,IL_) plays a more prominent role in matrix degradation compared to the rate of protease stimulation induced by mechanoinflammation.

Overall, these results represent proteolytic activity as a fundamental mechanism of post-injury cartilage degradation operating through distinct mechanoinflammatory and cytokine-driven inflammatory pathways. The framework may serve as an *in silico* platform for quantifying spatiotemporal protease activity in cartilage, which is difficult to resolve experimentally. Additionally, it may also support the virtual design of anti-inflammatory or protease-inhibiting therapeutic strategies for PTOA progression.

### 4.1 INJ-only and IL-only models

Simulations of injurious loading and IL-1 diffusion both produced substantial collagen and aggrecan loss by day 12, consistent with experimental observations. However, the spatial patterns of degradation differed between the two pathways (Figure 4&5).

For collagen, the INJ model agreed most with experiments in superficial and deep regions but underestimated the pronounced mid-depth collagen loss observed experimentally. This discrepancy suggests that mechanisms other than enzymatic proteolysis may be more prominent in this region, such as mechanical injury-induced fibril denaturation [54,55]. Excessive shear and tensile strains have been reported to cause fibril rupture and loss of collagen structural integrity independent of enzymatic cleavage [56–58], a process not explicitly incorporated in the current model formulation.

Conversely, the IL model reproduced collagen loss across most of the tissue depth, consistent with previous reports of bulk tissue degradation in response to inflammatory cytokines[26,50]. However, the model predicted greater collagen and aggrecan content loss in the superficial zone compared with experiment (Figure 4&5). This difference may arise from the boundary conditions imposed on IL-1 diffusion. Specifically, IL-1 was assumed to diffuse from the tissue surface with a constant diffusivity, which maintained high IL-1 concentration near the superficial region throughout the 12-day simulation period.

For aggrecan, model–experiment agreement was generally stronger in both INJ and IL conditions, suggesting that aggrecan loss might be more directly governed by proteolytic diffusion–reaction processes than collagen loss and is therefore better captured by the present framework. Although early aggrecan loss was experimentally observed on day 3, the model predicted only minimal aggrecan loss. However, the simulated profiles remained within the experimental 95% CI. This may reflect an early burst of aggrecan fragments out of the tissue in response to injury which is irrelevant to proteolytic-driven degradation [54]. In addition, the biological variability among the six experimental samples suggests that the observed day-3 differences between groups may partly reflect experimental uncertainty rather than a true limitation of the model.

### 4.2 Combined model INJ+IL

Simulations combining injurious loading and inflammation showed strong agreement with the experimental depth-dependent profiles on day 12 for both collagen and aggrecan.

Notably, the INJ+IL experiments did not demonstrate substantially greater collagen loss than INJ or IL alone relative to CTRL on day 12, whereas aggrecan loss was further aggravated, although still in a non-additive manner. The model captured this response through the interaction of the two catabolic pathways: IL-1-inflammation stimulation accelerated the transition of injury-induced catabolic cells to cell death, thereby limiting mechanoinflammation-driven protease secretion. This behavior is consistent with previous experimental observations showing that inflammatory cytokines exert a dual role in cartilage by both stimulating matrix-degrading protease expression and increasing cell death [50,59,60]. Within the present simulations, this dual action produced a transient increase in proteolytic activity shortly after injury, followed by a gradual shift toward IL-1-dominated degradation over longer timescales (Figure 4E & 5E). This is consistent with proteomics analysis reported by Wang et al. [60], which showed that inflammatory cytokines, such as IL-6 sustain MMP release to the culture medium for up to 21 days following combined injurious loading and IL-6 stimulation in human cartilage plugs [60]. Together, these findings support the view that mechanical injury-associated mechanoinflammation drives an acute proteolytic response, whereas IL-1 inflammatory signaling prolongs and sustains protease activity, ultimately governing longer-term matrix degradation. By resolving the distinct temporal dynamics of both pathways, the present framework provides a mechanistic basis for selecting and timing targeted therapeutic interventions.

Previous computational studies have also linked mechanical injury with catabolic cellular signaling. For example, Kapitanov et al. proposed a mathematical model in which mechanically induced cell damage initiates a catabolic response that evolves through the release of pro-inflammatory cytokines and reactive oxygen species [36]. The present framework shares similar mechanistic concepts, including protease-mediated degradation, and reaction-diffusion descriptions of the biochemical processes. However, our model extends these concepts by explicitly representing mechanoinflammation-driven and inflammatory cytokine-driven pathways as not only parallel but interacting processes. In most previous computational models, inflammatory signaling is treated primarily as a downstream consequence of the mechanical overloading[30,31,36]. In contrast, the present study represents pro-inflammatory cytokines as an independent driver of proteolytic activity alongside mechanoinflammatory injurious loading, enabling direct comparison of the individual and combined contributions of these mechanisms to cartilage degeneration.

### 4.3 Sensitivity analysis

Sensitivity analysis demonstrated that matrix degradation was influenced more by the amount of protease produced per catabolic cell than by the rate of protease stimulation in the INJ model (Figure 6A&B). This finding suggests that cartilage matrix loss is determined primarily by the local concentration of active proteases. In other words, rapid cellular activation alone is not sufficient to produce substantial matrix degradation if protease production per cell remains limited. Instead, both the abundance of catabolic cells, and severity of catabolic cell phenotype, reflected in protease production per cell, could be more important determinants of tissue damage than the rate at which cells are stimulated by mechanical overloading. This hypothesis can be tested in future studies by identifying distinct catabolic chondrocyte subpopulations and characterizing their respective protease expression profiles, for example using single-cell RNA sequencing[61].

In contrast to injury-induced stimulation, the rate of IL-1–driven protease stimulation had a larger effect on matrix degradation (Figure 6E&F). Two key factors explain this difference. First, IL-1 mediated stimulation acts on both healthy and catabolic cells, thereby involving a larger population of cells in protease secretion. Consequently, even moderate changes in the stimulation rate can lead to a considerable increase in cumulative protease production over time. Second, IL-1 stimulation is driven by diffusion of cytokines into the tissue, resulting in a sustained stimulus overtime. In contrast, injury-induced mechanoinflammatory stimulation is more acute and localized, reflecting the immediate cellular response to mechanical damage. As a result, increasing the rate of IL-1-driven stimulation amplifies the time-integrated protease production more strongly than equivalent changes in the injury-driven stimulus. These findings are consistent with previous experimental and computational studies demonstrating that pro-inflammatory cytokines sustain long-term catabolic activity in cartilage by continuously stimulating matrix-degrading enzymes [50,62].

### 4.4 Limitations

Some limitations in this work should be acknowledged. The model focuses on protease-mediated degradation driven by mechanoinflammatory injurious loading and IL-1 inflammatory cytokines, whereas other contributors to cartilage degeneration, such as oxidative stress, additional inflammatory cytokines (i.e., IL-6, and TNF-α), and protease inhibitors (i.e., tissue inhibitor of metalloproteinases, TIMPs), were not explicitly incorporated. Chondrocyte behavior was simplified into transitions between healthy, catabolic, and dead states governed by rate constants, thereby specific intracellular signaling pathways, spatial cell migration and the regulation of catabolic proteolytic activity were not incorporated in the current workflow. Although these processes could, in principle, be represented using specific intracellular signaling models integrated within the current framework [63], doing so would substantially increase the number of model parameters and associated uncertainty. In addition, several parameters (see Supplementary Material) were determined through model calibration and preliminary simulations rather than direct experimental calibration. Reducing uncertainty in these parameters will require further model validation using expanded experimental datasets, including gene expression and proteomic measurements of matrix degradation.

### 4.5 Conclusion

This study presents a mechanobiological computational framework integrating mechanoinflammation, IL-1 inflammatory signaling, and protease-driven matrix degradation to explain early cartilage degeneration following knee injury and the onset of PTOA. Our model was able to capture simultaneous aggrecan and collagen degeneration in injured and inflamed cartilage, providing a comprehensive mechanistic framework to analyze and separate different cartilage degeneration mechanisms. The framework could be used in the *in silico* design of therapeutic strategies and creation of virtual cohorts aimed at preventing long-term cartilage deterioration.

## Author contributions

**Moustafa Hamada:** Conceptualization, Data curation, Formal Analysis, Methodology, Visualization, and Writing – Original Draft Preparation. **Atte Eskelinen:** Conceptualization, Methodology, Writing – Review & Editing, Supervision, and Project administration. **Joonas Kosonen:** Conceptualization, Methodology, and Writing – Review & Editing. **Sanna Hakonen:** Conceptualization, Methodology, and Writing – Review & Editing. **Cristina Florea:** Conceptualization, Methodology, and Writing – Review & Editing. **Alan Grodzinsky:** Conceptualization, Methodology, Resources, and Writing – Review & Editing. **Rami Korhonen:** Conceptualization, Methodology, Writing – Review & Editing, Supervision, Project administration, and Funding acquisition. **Petri Tanska:** Conceptualization, Methodology, Writing – Review & Editing, Supervision, Project administration, and Funding acquisition.

