## Supporting information for "A computational framework to model cartilage degeneration induced by mechanoinflammation and cytokine-driven inflammation in post-traumatic osteoarthritis"

Telephone number: **+358449194157, +358294453164**

### **S1 Parameters in the biomechanical model**

In the biomechanical model, cartilage was modeled as a fibril-reinforced porohyperelastic material with Donnon osmotic swelling. Mechanical properties of the tissue are considered through a non-fibrillar matrix (representing aggrecan) and fibrillar matrix (collagen fibrils). The non-fibrillar part is described with Cauchy stress tensor of a neo-Hookean solid material. And it is formulated as follows:

| $\boldsymbol{\sigma}_{\mathrm{nf}}=K_{\mathrm{nf}}\frac{\ln\left( J \right)}{J}\mathbf{I}+\frac{G_{\mathrm{nf}}}{J}\left( \mathbf{F}\cdot\mathbf{F}^{T}-J^{\frac{2}{3}} \mathbf{I} \right),$ | (S.E 1) |
| --- | --- |

where $K_{\mathrm{nf}}$ is bulk modulus, $G_{\mathrm{nf}}$ is shear modulus, $\mathbf{F}$ is deformation gradient tensor, $J$ is volumetric deformation and $\mathbf{I}$ is unit tensor.

$K_{\mathrm{nf}}$ and $G_{\mathrm{nf}}$ is defined respectively as follows:

|  | $K_{\mathrm{nf}}=\frac{E_{\mathrm{nf}}}{3\left( 1-2\nu_{\mathrm{nf}} \right)} ,$ | (S.E 2) |
| --- | --- | --- |

|  | $G_{\mathrm{nf}}=\frac{E_{\mathrm{nf}}}{2\left( 1+\nu_{\mathrm{nf}} \right)},$ | (S.E 3) |
| --- | --- | --- |

where $E_{\mathrm{nf}}$ is the Young’s modulus, and $\nu_{\mathrm{nf}}$ is the nonfibrillar matrix Poisson’s ratio.

For the fibrillar matrix, the stress was modeled with a linear elastic material:

| $\sigma_{f}=\left\{ \begin{aligned} E_{f}\varepsilon_{f}, &if \varepsilon_{f}\geq0, \\ 0, &\mathrm{if} \varepsilon_{f}<0, \end{aligned} \right.$ | (S.E 4) |
| --- | --- |

where $E_{f}$ is the fibril modulus and $\varepsilon_{f}$ is the logarithmic fibril strain. $\varepsilon_{f}$ is described as:

|  | $\varepsilon_{f}$ = ln (\|\|**F *e***_f,0_\|\|) | (S.E 5) |
| --- | --- | --- |

where ***e***_f,0_ is the unit vector of the initial fibril orientation.

A fibril consists of 2 primary fibrils that define the curved main structure of young bovine cartilage and 7 secondary fibrils to define the randomly oriented fibrils. Therefore, for fibril *j*, the Cauchy stress tensor is:

| ${\boldsymbol{\sigma}_{f}}^{j}=\left\{ \begin{aligned} c_{col,0}C\sigma_{f}\boldsymbol{e}_{f}\otimes\boldsymbol{e}_{f}, &for primary fibrils, \\ c_{col,0}\sigma_{f}\boldsymbol{e}_{f}\otimes\boldsymbol{e}_{f}, &for secondary fibrils, \end{aligned} \right.$ | (S.E 6) |
| --- | --- |

where $c_{col,0}$ is the depth-dependent collagen fraction per solid volume, $C$ is the density ratio between primary and secondary fibrils, and $\boldsymbol{e}_{f}$ is the current normalized fibril orientation vector.

The fluid flow is described according to Darcy’s law:

|  | $q=-k\nabla p,$ | (S.E 7) |
| --- | --- | --- |

where $q$ is the flow rate, $k$ is the permeability, and $\nabla p$ is fluid pressure gradient.

The Donnan osmotic pressure gradient is described in equilibrium as following:

|  | $\Delta\pi=\phi_{\mathrm{int}}\mathrm{RT}\left( \sqrt{c_{F}^{2}+4\frac{\left( \gamma_{\mathrm{ext}}^{\pm} \right)^{2}}{\left( \gamma_{\mathrm{int}}^{\pm} \right)^{2}}c_{\mathrm{ext}}^{2}} \right)-2\phi_{\mathrm{ext}}\mathrm{RT}c_{\mathrm{ext}},$ | (S.E 8) |
| --- | --- | --- |

where $C_{F}$ is the depth-dependent fixed charge density concentration, $\phi_{\mathrm{int}}$, $\phi_{\mathrm{ext}}$, $\gamma_{\mathrm{int}}^{\pm}$, and $\gamma_{\mathrm{ext}}^{\pm}$ are internal and external osmotic coefficients and internal and external activity coefficients, respectively. $c_{\mathrm{ext}}$ is the external salt concentration, $R$ is the molar gas constant and $T$ is the absolute temperature.

The chemical expansion stress is

|  | $T_{c}=a_{0}c_{F}\exp\left( -\kappa\frac{\gamma_{\mathrm{ext}}^{\pm}}{\gamma_{\mathrm{int}}^{\pm}}\sqrt{c^{-} \left( c^{-}+c_{F} \right)} \right),$ | (S.E 9) |
| --- | --- | --- |

where $a_{0}$, $\kappa$ are material constants and $c^{-}$ is the anion concentration.

Finally, the total stress tensor comprises the fibril matrix stress, the non-fibril matrix stress, the fluid pressure gradient, and chemical expansion:

|  | $\boldsymbol{\sigma}_{\mathrm{tot}}\boldsymbol{=}\sum_{j=1}^{\mathrm{totf}} {\boldsymbol{\sigma}_{f}}^{j}+\boldsymbol{\sigma}_{\mathrm{nf}}\mathbf{-}\Delta\pi\mathbf{I-}T_{c}\mathbf{I-}\mu_{f}\mathbf{I},$ | (S.E 10) |
| --- | --- | --- |

Where $\mu_{f}$ is the chemical potential, and $\mathrm{totf}$ represent the sum of primary and secondary fibrils.

The main material parameters for the material model are listed in Table S1.

**Table S1. Material model parameters.** z represents the normalized depth of the tissue from the surface (surface = 0, bottom = 1).

| Material parameter | Value | Description | Reference |
| --- | --- | --- | --- |
| E_f_ (MPa) | 20.0 | Fibril network modulus | ^1^ |
| E_nf_ (MPa) | 0.16 | Non-fibrillar matrix modulus | ^2^ |
| ν_nf (-)_ | 0.4 | Poisson’s ratio of the non-fibrillar matrix | ^3^ |
| k  (10^-15^ m^4^/(N⋅s)) | 1.3 | Hydraulic permeability | ^2^ |
| C [-] | 3.009 | Density ratio between primary and secondary collagen fibrils | ^2^ |
| Distributions | Value | Description | Reference |
| $n_{\text{f,0}}$ | $0.85-0.1z$ | Initial depth-wise fluid fraction | ^1^ |
| $c_{col,0}$ | $-187.1z^{6}+328.6z^{5}+22.9z^{4}-$  $363.4z^{3}+247.8z^{2}-47.9z+8.782$ | Initial depth-wise collagen fraction | ^4^ |
| $c_{F,0}$ (mEq · ml^-1^) | $-4.4z^{6}+15.2z^{5}-21.0z^{4}+14.9z^{3}$  $-5.8z^{2}+1.1z+0.03$ | Initial depth-wise fixed charge density | ^1^ |

### **S2 Parameters in the biochemical model**

The simulated cartilage degradation consisted of the loss of both collagen and aggrecan content^5^. Both components were degraded via cell-driven proteolytic activity: in response to injurious loading and IL-1 diffusion^6^. Healthy and catabolic cells exhibit increased expression of proteases such as matrix metalloproteinases (MMP) and aggrecanases due to mechanoinflammation and cytokine-driven inflammation. MMPs are primarily responsible for the collagen fibrils degradation, while aggrecanases are linked with aggrecan loss^7–10^. Changes in MMP ($C_{mmp, total}$) and intact collagen ($C_{\mathrm{col}}$) concentrations are described in Equations (10 &12) in the main text. The degraded collagen concentration is given by:

| $\frac{\partial C_{\mathrm{cold}}}{\partial t}=f_{agg,protect}\left( C_{\mathrm{agg}} \right)k_{mmp,catalytic} C_{mmp, total}\left( \frac{C_{\mathrm{col}}}{C_{\mathrm{col}}+K_{m,mmp}} \right),$ | (S.E 11) |
| --- | --- |

where $k_{mmp,catalytic}$ is the catalytic activity rate of MMPs, while $K_{m,mmp}$ is the Michaelis constant describing the MMP concentration at which collagen cleavage proceeds at half-maximum rate.

$f_{agg,protect}\left( C_{\mathrm{agg}} \right)$ represents modulation of collagen loss by the local aggrecan content and it is defined as following:

| $f_{agg,protect}\left( C_{\mathrm{agg}} \right)=\frac{1}{1+\left( \frac{C_{\mathrm{agg}}+C_{\mathrm{aggd}}}{k_{\mathrm{act}}} \right)^{-n}},$ | (S.E 12) |
| --- | --- |

where $C_{\mathrm{agg}}$ and $C_{\mathrm{aggd}}$ are the concentrations of intact and degraded aggrecan respectively, $k_{\mathrm{act}}$ defines aggrecan concentration at half-maximal MMP activity, and $n$ is the Hill coefficient of MMP activity.

The change in aggrecanase ($C_{aga,tot}$) and intact aggrecan ($C_{\mathrm{agg}}$) concentrations are defined in equations (11 & 13) in the main text. The biosynthesis ($R_{\mathrm{biosynthesis}}$) of aggrecan is described below:

| $R_{\mathrm{biosynthesis}}=P_{\mathrm{agg}}\left( 1+\frac{0.9(1-z)}{H} \right),$ | (S.E 13) |
| --- | --- |

where $P_{\mathrm{agg}}$ is the basal amount of aggrecan produced by the healthy cells, $z$ is the normalized axial co-ordinate within the tissue (z = 0 at the surface, z = 1 at the bottom), and $H$ is the thickness of the tissue.

The change in degraded aggrecan concentration $C_{\mathrm{aggd}}$ in the tissue is described as:

| $\frac{\partial C_{\mathrm{aggd}}}{\partial t}=D_{\mathrm{aggd}}\nabla^{2}C_{\mathrm{aggd}}+k_{aga,catalytic}C_{aga,total}\left( \frac{C_{\mathrm{agg}}}{C_{\mathrm{agg}}+K_{m,aga}} \right),$ | (S.E 14) |
| --- | --- |

The values used for the model parameters are detailed in Table S2.

**Table S2. Biochemical model parameters**

| Model parameter | Value | Description | Reference |
| --- | --- | --- | --- |
| $C_{cell,h} (cells.m^{-3})$ | 1.5 × 10^14^ | Initial healthy cell concentration **(Eq 2)** | ^10^ |
| $k_{\mathrm{INJ}} (-)$ | 0.35 | Maximum allowed cell damage **(Eq 7)** | ^6^ |
| $k_{\mathrm{mmp}}$, $k_{\mathrm{aga}}$  (10^-21^ mol) | 0.25 | Production of MMPs and aggrecanases from damaged cells **(Eq 8)** | Model fit |
| $k_{rate, mmp}$, $k_{rate,aga}$  (${10}^{-5}s^{-1}$) | 3.6 | MMPs and aggrecanases production rate **(Eq 10 & 11)** | Model fit, ^10^ |
| $k_{mmp,cold}\left( m^{3}{\cdot mol}^{-1}\cdot s^{-1} \right)$ | 4.7 ×10^-4^ | Rate of MMP binding to degraded collagen **(Eq 10)** | ^10^ |
| $n_{R,cold}(-)$ | 320 | Number of MMP binding sites on degraded collagen **(Eq 10)** | ^10^ |
| $k_{\mathrm{act}} ({10}^{-4}mol\cdot m^{-3})$ | 3 | Aggrecan concentration at half maximum of MMP activity **(S.E 12)** | ^10^ |
| $n (-)$ | 6 | Hill coefficient of MMP activity **(S.E 12)** | ^10^ |
| $k_{mmp,catalytic} {(s}^{-1})$ | 1.5 | Catalytic activity rate of MMPs with collagen fibrils **(Eq 12)** | ^10^ |
| $K_{m,mmp} (mol\cdot m^{-3})$ | 0.0021 | Michaelis constant for MMPs **(Eq 12)** | ^10^ |
| $k_{loss,aga}({10}^{-4}s^{-1})$ | 1 | Aggrecanase degradation rate **(Eq 11)** | ^10^ |
| $P_{\mathrm{agg}}({10}^{-22}\mathrm{mol}s^{-1})$ | 2.4 | Basal amount of aggrecan production **(S.E 13)** | ^10^ |
| $C_{\mathrm{tar}}(mol\cdot m^{-3})$ | 0.011635 | Target aggrecan concentration **(Eq 13)** | Model fit, ^10^ |
| $k_{aga,catalytic}{(s}^{-1})$ | 0.9 | Catalytic activity rate of aggrecanases with aggrecan **(Eq 13)** | ^10^ |
| $K_{m,aga}({10}^{-5}mol\cdot m^{-3})$ | 5 | Michaelis constant for aggrecanases **(Eq 13)** | ^10^ |
